# Heterogeneous Tfh cell populations that develop during enteric helminth infection predict the quality of type 2 protective response

**DOI:** 10.1101/2021.10.20.465079

**Authors:** Aidil Zaini, Lennard Dalit, Amania A. Sheikh, Daniel Thiele, Yan Zhang, Jessica Runting, Grace Rodrigues, Judy Ng, Michael Bramhall, Sebastian Scheer, Lauren Hailes, Joanna R. Groom, Kim L. Good-Jacobson, Colby Zaph

## Abstract

T follicular helper (Tfh) cells are an important component of the germinal centre (GC)-mediated humoral immunity. Yet, how regulation of Tfh-GC responses impacts on effective responses to helminth infection are poorly understood. Here we show that chronic helminth *Trichuris muris* infection fails to induce Tfh-GC B cell responses, with Tfh cells expressing T-bet and IFN-γ. In contrast, Tfh cells that express GATA-3 and IL-4 dominate responses to an acute, resolving infection. Accordingly, heightened expression and increased chromatin accessibility of Th1- and Th2 cell-associated genes is observed in chronic and acute induced Tfh cells, respectively. However, both acute and chronic Tfh cell populations retained the capacity to produce IL-21 in spite of the Th-biased response. Blockade of Tfh-GC interactions impaired type 2 immunity, highlighting the protective role of GC-dependent Th2-like Tfh cell responses against helminths. Collectively, these results provide new insights into the protective roles of Tfh-GC responses and identify distinct transcriptional and epigenetic features of Tfh cells that emerge during resolving or chronic helminth infections.

**Author summary:** About a quarter of the world population is afflicted with parasitic worm infection. Although deworming drugs can reduce the levels of the infection, they fail to prevent reinfections. Therefore, the most sustainable goal is to develop vaccines against human helminth parasites, which has been extremely challenging due to the lack of understanding of host-parasite interactions. While the protective roles of T helper 2 (Th2) cells are well established, the regulation of T follicular helper (Tfh) cells and their roles during helminth infection remain poorly defined. In this study, we describe the differential regulation of Tfh cell responses during chronic non-protective vs acute protective responses during helminth infection. We show that Tfh cells during chronic infection are rare and have strikingly different characteristics to acute-induced Tfh cells, which appear to be more like Th2 cells. Specifically, we show that blockade of Th2-like Tfh cell response during acute infection results in the host failing to expel the worms. Our study identifies that Tfh cell populations that emerge during chronic and acute infection are strikingly heterogeneous and critically important in mediating protective immune responses against helminths.

## Introduction

The germinal center (GC) reaction is a highly dynamic process through which high-affinity and long-lived antibodies and memory B cells develop in response to infections, and is often used as the basis of vaccinations, primarily against viral pathogens. However, there are currently no viable human vaccines against helminth parasites that afflict billions of people world-wide [1]. The lack of helminth vaccines may reflect the complexity and diversity of helminths, the biology of the infectious cycle, as well as the lack of knowledge on the precise requirements for immunity to infection. Although Th2 cells are the key to protective responses against helminths, the role of GCs in promoting effective responses to acute helminth infection, or whether their dysregulation contributes to ineffective clearance during chronic infection, remains poorly defined [2].

CXCR5-expressing T follicular helper (Tfh) cells aid B cell responses through the production of cytokines and direct contact-mediated interactions such as CD40-CD40L [3]. Tfh cell differentiation is dependent upon Bcl6, the Tfh cell lineage-defining transcription factor (TF). Mechanistically, Bcl6 represses non-Tfh cell lineage genes, including Th1 cell-defining T-bet and Th2 cell-associated Gata3 [4–6], as well as through the repression-of-repressors mechanism [7,8]. Despite Bcl6-dependent repression of T-bet and Gata3, distinct subpopulations of Tfh cells that share Th1 and Th2 phenotypes develop during different types of infection [9,10]. In response to viral infection, Tfh cells initially co-express Bcl6 and T-bet and the interplay between these TFs plays an important role in the establishment of Tfh versus Th1 cell lineage [11,12] in a context-dependent manner [13–15]. Type 2 immune stimuli such as allergens and helminths promote Tfh cells that mainly produce IL-4, a cytokine that is required for coordinating the overall type 2 immune response, as well as IgG1 and IgE class switching [16–18], as well as for coordinating the overall type 2 immune response. The diverse Tfh cell phenotypes suggest that tailoring of Tfh cells is central to the development of protective CD4^+^ T cell responses across pathogens [9,10]. However, how the type of the immune response (type 1 vs type 2) against one class of pathogen regulates the flexibility and functionality of Tfh cell responses remains unclear.

In the current study, we characterized Tfh cell and GC B cell responses in the draining mesenteric lymph nodes (mLNs) during acute high dose (HD) and chronic low dose (LD) *Trichuris muris* (*T. muris*) infection. The *T. muris* model allows us to delineate Tfh-GC responses in either Th1- or Th2-biased responses against a single pathogen [19]. Our characterization revealed that acute but not a chronic *T. muris* infection drives the development of a robust Tfh cell response and GC formation. Tfh cells from HD infected mice displayed hallmarks of Th2 cells transcriptionally and epigenetically. Under Th2 polarizing conditions, *ex vivo* HD-derived Tfh cells maintained canonical Tfh cell markers, including IL-4 and IL-21 expression. LD infection promoted fewer Tfh cells and a poorer GC response when compared to HD infection, and these Tfh cells exhibited Th1 cell attributes. Despite differences in Th1/Th2 cell phenotypes, both HD- and LD-induced Tfh cells remained expressing IL-21. Blockade of the Th1 cell response by T cell-intrinsic T-bet deletion resulted in a significant increase in Tfh cells during chronic infection, further demonstrating a relationship between a robust Tfh cell response and immunity to helminth infection. Finally, blockade of GC responses during HD infection promoted susceptibility to the parasite. Taken together, our results show that Tfh cell development is differentially regulated during acute and chronic helminth infection. Importantly, Tfh cell response shapes the quality of type 2 immunity, which can be further therapeutically manipulated to control chronic helminth infection.

## Results

### Acute high-dose helminth infection drives a potent Tfh-GC response

Although both type 1 and type 2 immune stimuli drive Tfh cell development, the functional capacities of Tfh1 and Tfh2 cells are discrete. In turn, distinct Tfh cell function regulates the outcome of humoral immune responses and the host’s immunity [13,18,20]. The distinct functions of different subpopulations of Tfh cells suggest that they play key non-redundant roles in regulating immunity to a wide range of immune stimuli. To first characterize Tfh and GC B cell regulation during helminth-induced Th1 and Th2 responses, we infected C57BL/6J mice with the helminth parasite *T. muris* in both chronic (LD) and acute (HD) settings and evaluate how these responses correlate with protection at different time points post infection (pi) (**Fig 1A**). Consistent with previous studies [21,22], worm burden in HD-infected mice progressively decreased from 7 to 14 days pi, with almost all worms cleared by day 21 pi. (**Fig 1B**). In contrast to HD infection, LD-infected mice failed to clear the parasite, maintaining a consistent and chronic worm burden to day 35 pi. (**Fig 1B**). These results indicate that C57BL/6J mice were capable of effectively clearing worms post-HD infection, whereas LD infection results in susceptibility to the parasite.

**Fig 1.**
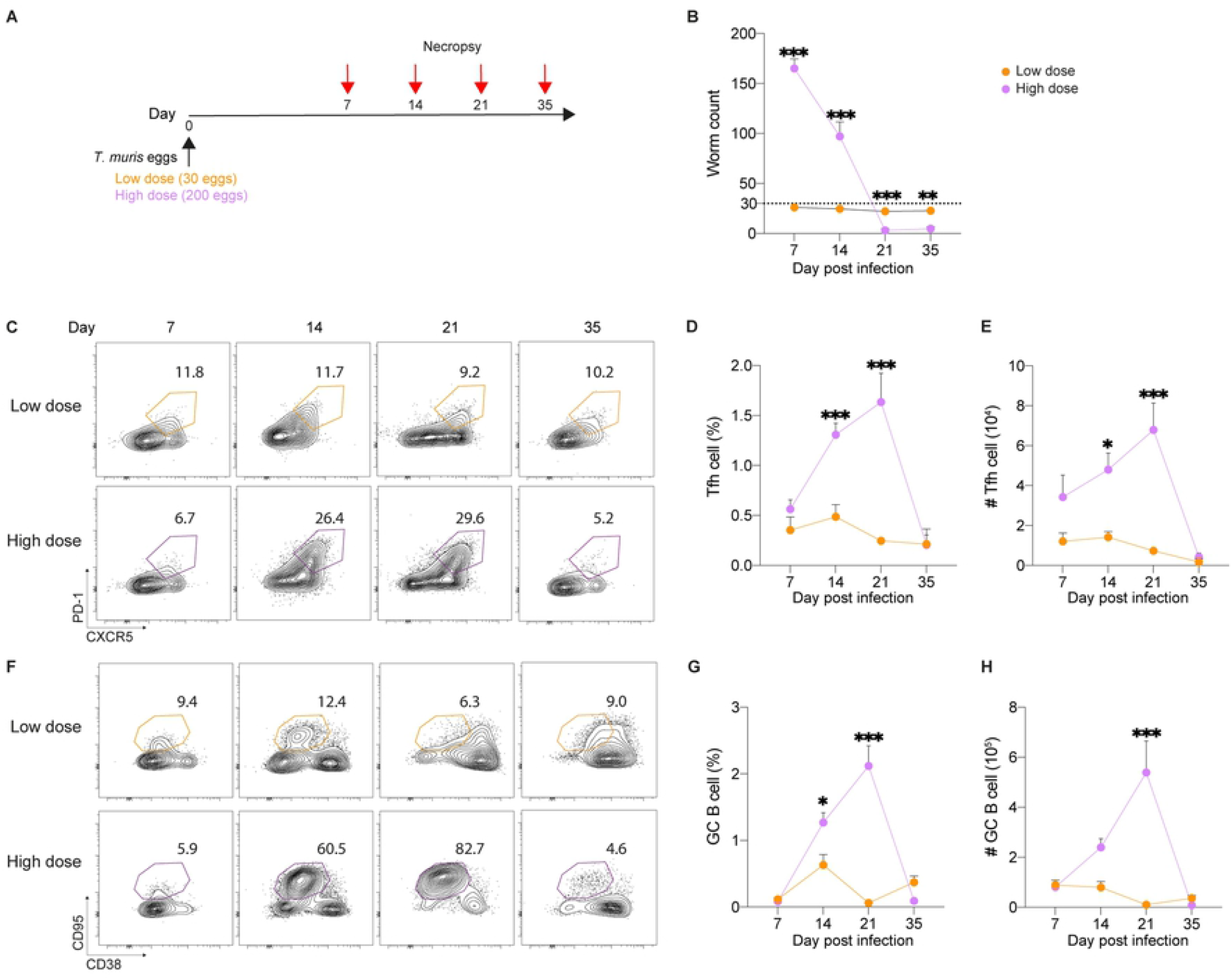
Acute but not chronic helminth infection drives Tfh cell development and GC reactions. (A) C57BL/6J mice were infected with acute (high dose) and chronic (low dose) *T. muris*, and caecum and mLNs were analyzed at d7, d14, d21 and d35 pi. (B) Worm burden analysis in the caecum at indicated days pi. (C) Representative flow cytometric plots showing frequency of Tfh cells (CD4^+^ Bcl6^+^ CXCR5^+^ PD-1^+^) at indicated days pi. (D and E) Frequency (E) and total numbers of (E) of Tfh cells. (F) Representative flow cytometric plots showing frequency of GC B cells (CD19^+^ IgD^-^ CD95^+^ CD38^-^) at indicated days pi. (G and H) Frequency (G) and total numbers of (H) of GC B cells. Data shown are representative of three independent experiments (n = 3–6 mice per group). Statistical significance was determined with a one-way ANOVA. Error bars represent means ± SEM. *p<0.05, **p<0.01, ***p<0.001. A dotted line in (A) represents the levels of low dose infection.

We next characterised the Tfh/GC response during non-protective and protective responses using LD- and HD-infection models. We used the canonical Tfh cell (CD4^+^ BCL6^+^ CXCR5^+^ PD-1^+^) and GC B cell (B220^+^ IgD^-^ CD95^hi^ CD38^+^) markers to identify Tfh and GC B cells in the mLNs at day 7, 14, 21 and 35 following LD or HD infection. Flow cytometric analysis revealed the frequencies and numbers of HD-induced Tfh cells peaked at d21 pi (**Fig 1C-1E**). In contrast, Tfh cell frequencies and numbers were lower during LD infection at all timepoints (**Fig 1C-1E**). The GC B cell response paralleled the Tfh cell response during LD and HD infection (**Fig 1F-1H**). We found significantly fewer GCs in the mLNs during LD infection (**Figs 2A-2C and S1A**). Further, the size of the GCs as determined by the GL7^+^/IgD^+^ area was also significantly smaller in the LD-infected mLNs (**Fig 2D**), suggesting that LD infection fails to induce GC B cell expansion. Consistent with the development of a protective Th2 cell response in resistant mice [23], we found high levels of serum parasite-specific IgG1 and low levels of IgG2c (**Fig 2E and 2F**). Surprisingly, despite a poor Tfh-GC response, LD-infected mice were capable of producing class-switched IgG2c (**Fig 2E and 2F**). Taken together, our results show that potent Tfh-GC responses correlate with type 2-mediated protection following HD *T. muris* infection.

**Fig 2.**
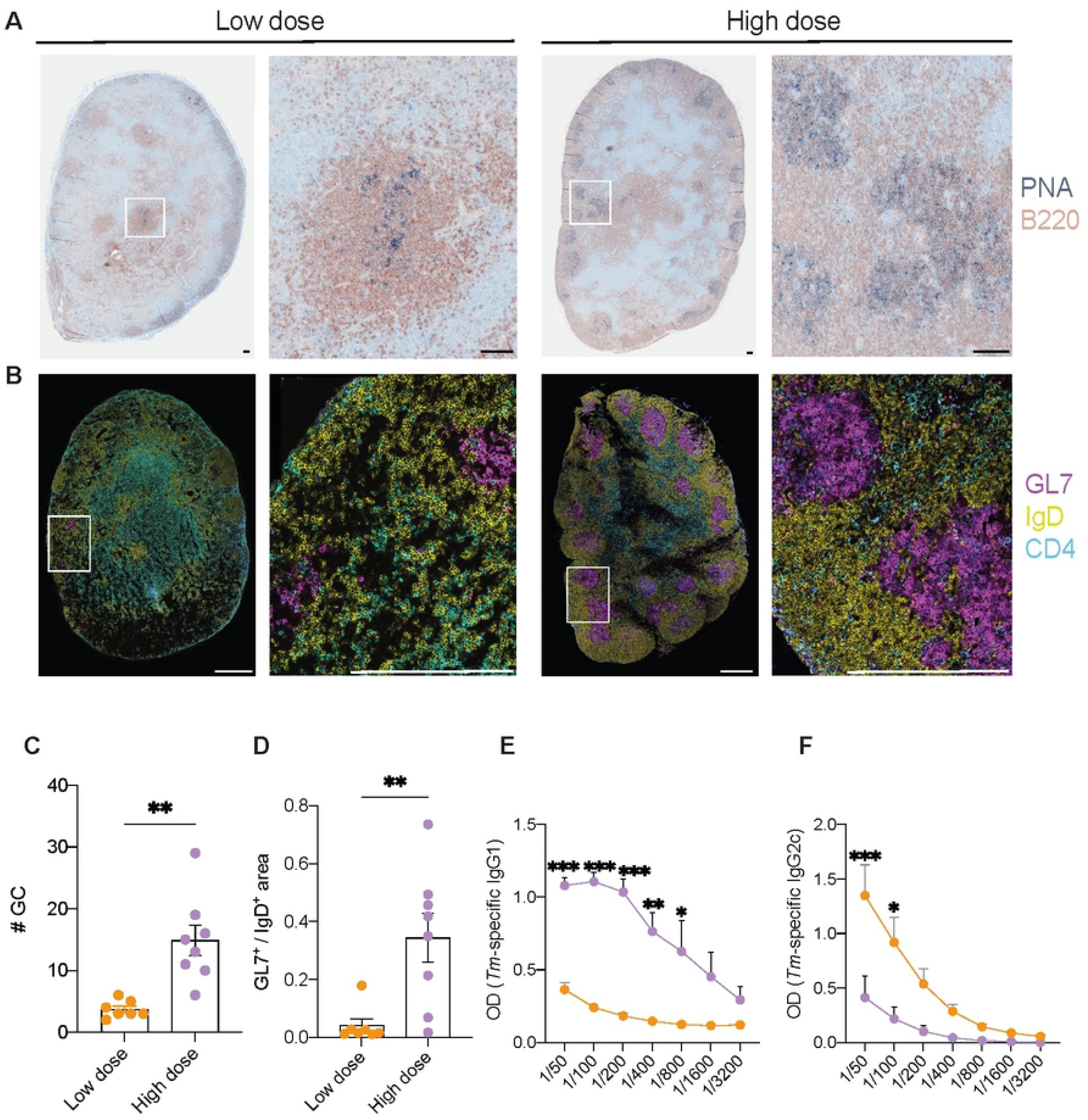
Robust GC formation in acute helminth infection*-*induced draining mLNs. C57BL/6J mice were infected with acute (high dose) and chronic (low dose) *T. muris*, and mLNs were analyzed at d21 pi. (A and B) Representative histological analyses of mLN tissue: B220 (red) and PNA (blue). Scale bars, 100 μm. (B) Representative confocal micrographs of mLN tissue: GL7 (magenta), IgD (yellow) and CD4 (cyan). Scale bars, 100 μm. (C and D) Quantitation of total numbers of GC/mLN section (GL7^+^ IgD^-^) (C) and GC size (GL7^+^/IgD^+^ area). (E and F) *T. muris*-specific serum IgG1 (E) and IgG2c (F). Data shown in (E and F) are representative of three independent experiments (n = 3–6 mice per group). Statistical significance was determined with a two-tailed Student’s t test. Error bars represent means ± SEM. *p<0.05, **p<0.01, ***p<0.001. *Tm*; *Trichuris muris*.

### Kinetics of IL-4 and IL-21 expression during Th1- and Th2-skewed helminth infection

As we had observed robust GC responses that developed progressively throughout HD but not LD *T. muris* infection, we next generated IL-4-AmCyan/IL-13-DsRed/IL-21-GFP (IL-4-13-21) triple reporter mice [24–26] to dissect the temporal expression of these cytokines in Tfh cells. First, we failed to detect any IL-13-DsRed expression in both CD4^+^ CD44^hi^ CXCR5^-^ PD-1^-^ non-Tfh cells and Tfh cells at d7-21 post-LD- and -HD *T. muris* infection (**S1B Fig**). This finding complement previous studies showing that IL-13 expression is typically expressed in CD4^+^ T cells in the non-lymphoid tissue such as lungs but not draining lymph nodes (LNs) [27,28]. In addition, the absence of IL-13-producing Tfh cells in response to *T. muris* infection is consistent with results from infections with the helminth *Nippostrongylus brasiliensis* (*N. brasiliensis*) [29]. The expression of IL-4 single-producing Tfh cells in both LD and HD infections peaked at d14, with the higher levels of single IL-4-expressing Tfh cells observed in HD than that of LD infection (**Fig 3A and 3B**). Consistent with a population of Tfh cells that co-produce both IL-4 and IL-21 in response to *N. brasiliensis* infection [18], we also observed Tfh cells that co-expressed IL-4 and IL-21 in both LD and HD *T. muris* infections (**Fig 3A-3C**). Similar to single IL-4-producing Tfh cells, both LD and HD infection drove comparable levels of IL-4 and IL-21 double-positive Tfh cell frequencies prior to the peak of Tfh-GC responses (i.e., d7 and d14 pi) (**Fig 3A-3C**). Unlike single IL-4 expression that peaked at d14, the expansion of single IL-21 expression in both LD- and HD-induced Tfh cells dropped at this time point and both infection doses consistently drove a comparable proportion of IL-21 expression in Tfh cells throughout the course of infection (**Fig 3A and 3D**). Collectively, these results indicate that HD-but not LD-induced Tfh cells are potent producers of IL-4, yet LD-induced cells remained able to produce similar levels of IL-21 during HD infection.

**Fig 3.**
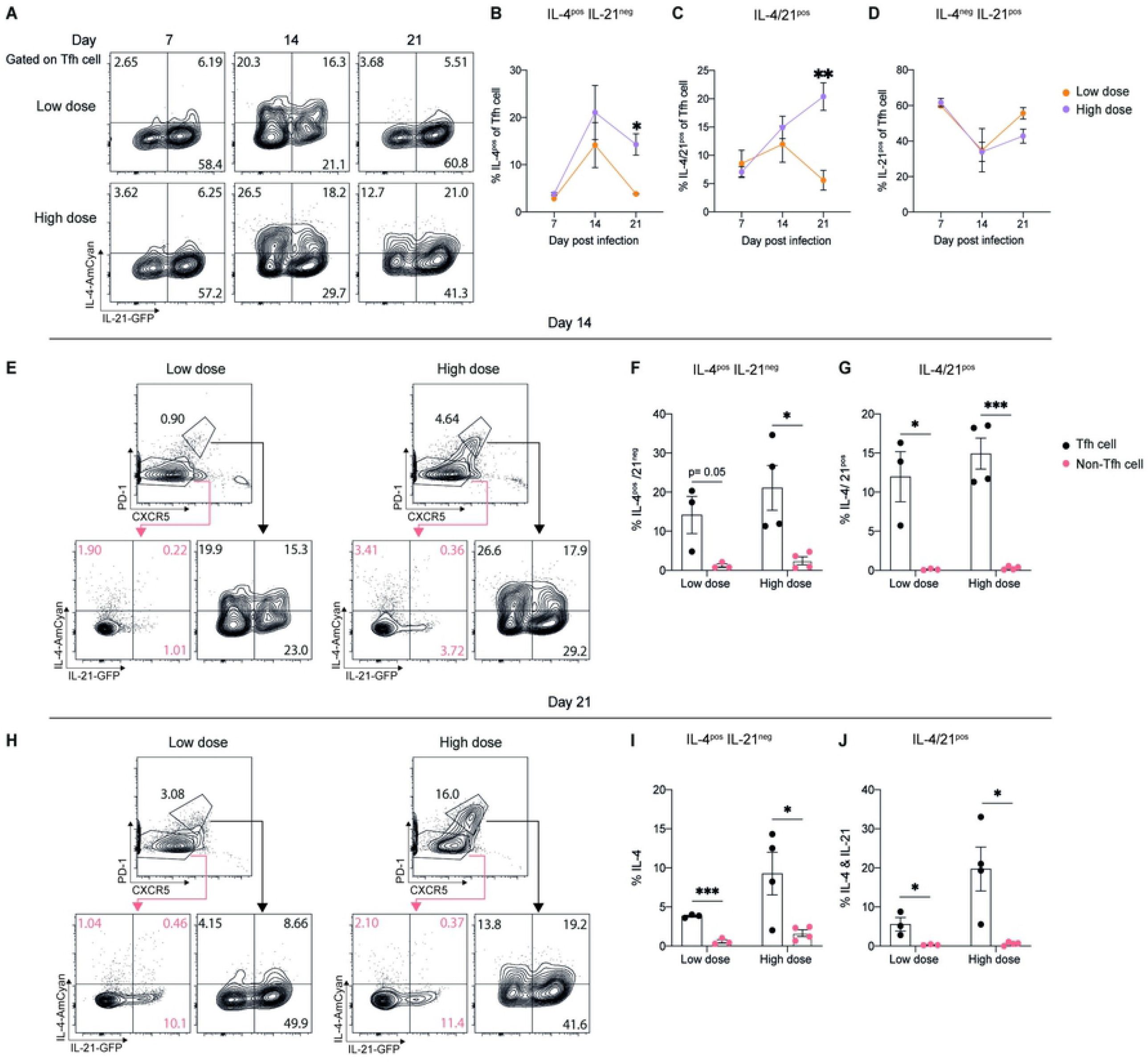
Kinetics of IL-4 and IL-21 expression in Tfh cells during acute and chronic helminth infection. IL-4-AmCyan/IL-13-DsRed/IL-21-GFP (IL-4-13-21) triple reporter mice were infected with acute (high dose) and chronic (low dose) *T. muris*, and mLNs were analyzed at d7, d14, and d21 pi. (A) Representative flow cytometric plots showing frequency of indicated populations for expression of IL-4 and IL-21 gated on Tfh cells (CD4^+^ CD44^hi^ CXCR5^+^ PD-1^+^) at indicated days pi. (B-D) Frequency of IL-4^pos^ IL-21^neg^ (B), IL-4/21^pos^ (B) and IL-4^neg^ IL-21^pos^ (C) populations of Tfh cells at indicated days pi. (E and H) Representative flow cytometry plots showing frequency of indicated populations for expression of IL-4 and IL-21 gated on Tfh cells (CD4^+^ CD44^hi^ CXCR5^+^ PD-1^+^) and non-Tfh cells (CD4^+^ CD44^hi^ CXCR5^-^ PD-1^-^) at d14 (E) and d21 (H) pi.(F-J) Frequency of IL-4^pos^ IL-21^neg^ (F and I) and IL-4/21^pos^ (G and J) populations of Tfh and non-Tfh cells at d14 (F and G) and d21 (I and J) pi. Data shown in are representative of three independent experiments (n = 3–4 mice per group). Statistical significance was determined with a two-tailed Student’s t test. Error bars represent means ± SEM. *p<0.05, **p<0.01, ***p<0.001.

Multiple reports demonstrate that Tfh cells are major producers of IL-4 in draining LNs in response to type 2 inflammatory stimuli [16,17,24]. Given HD *T. muris* infection potently drives IL-4-mediated Th2 cell responses [30], we sought to further clarify the producers of IL-4 during *T. muris* infection. At d14 and d21 pi, we observed that LD- and HD-induced Tfh cells were indeed the major producers of IL-4 with a substantial proportion that co-expressed IL-21 when compared to non-Tfh cell populations (**Fig 3E-3J**). Interestingly, non-Tfh cell populations also expressed IL-21 at d14 and the levels of IL-21 expression in these cells were more prominent at d21 pi (**Fig 3E and 3H**). Thus, IL-21 production was not expressed exclusively by Tfh cells. Taken together, these data suggest that Tfh cells are major producers of IL-4, a proportion of which is co-expressed with IL-21.

### B cell-intrinsic IL-4Rα signals are dispensable for immunity to helminths

Given we demonstrated that HD-induced Tfh cells expressed higher amounts of IL-4 at the peak of GC reactions (i.e., d21 pi) when compared to LD infection, we next sought to determine the effects of IL-4Rα signals on the development of GC responses during HD *T. muris* infection. Initially, we infected global IL-4Rα knockout mice (IL-4Rα^-/-^) with HD *T. muris* eggs, which harboured a higher worm burden than control (**S2A Fig**). Despite increased Th1 cell frequencies and reciprocal reduction in the proportion of Th2 cells (**S2B and S2C Fig**), we found no difference in Tfh cell frequencies. This confirms a previously published work that unlike Th2 cells, Tfh cell development does not require IL-4/IL-4Rα signals [31]. However, unlike Tfh cells, GC B cell frequencies were reduced in IL-4Rα^-/-^ mice (**S2E Fig**). Consistent with the requirement of IL-4 in class-switching of IgG1 during type 2 responses [16,18], we further showed that GC B cells in infected IL-4Rα^-/-^ mice had almost absent IgG1 expression (**S2F Fig**). However, these observations could not discern the potential protective B cell-intrinsic roles of IL-4Rα signals. Therefore, we generated bone marrow (BM) chimeric mice lacking IL-4Rα in their B cells only (B-IL-4Rα^-/-^), whereby BM cells from IL-4Rα^-/-^ and μMT mice were mixed in 1:4 ratio to reconstitute lethally irradiated recipients (**S2G Fig)**. The loss of IL-4Rα expression on B cells indicates the success of B-IL-4Rα^-/-^ chimera generation (**S2H Fig**). In contrast to IL-4Rα^-/-^ mice, we found that B-IL-4Rα^-/-^ mice were resistant to HD infection, although worm expulsion was slightly delayed at d14 and 21 pi (**S2I Fig**). As expected, we did not observe any changes in Tfh cell frequencies (**S2J Fig**). However, B-IL-4Rα^-/-^ mice had reduced GC B cell frequencies at d14 and 35, but unexpectedly, not at d21 pi (**S2K Fig**). Of note, B-IL-4Rα^-/-^ chimeric mice had a delayed GC response (**S2K Fig**) when compared to C57BL/6J mice (**Fig 1G-1H**). Consistent with IL-4Rα^-/-^ mice (**S2F Fig**), IgG1-switched GC B cells decreased (**S2L Fig**), albeit not to the same extent in IL-4Rα^-/-^ mice at d21 pi. Parasite-specific IgG1 (**S2M-S2O Fig**) but not IgG2c (**S2P-S2R Fig**) synthesis was also impaired in B-IL-4Rα^-/-^ mice at all time points observed. Thus, although IL-4Rα in B cells is required for IgG1 synthesis and GC B cell generation, IL-4Rα expression on B cells is dispensable for type 2 protective responses.

### Helminth infection-induced Tfh cells are transcriptionally diverse

To begin to identify the mechanisms controlling the differential Tfh cell/GC responses following LD and HD *T. muris* infection, we examined the transcriptional profile of CD4^+^ CD44^hi^ FOXP3^-^ Ly6C^-^ CD162^-^ PD-1^+^ CXCR5^+^ Tfh cells isolated from the mLN of HD- and LD-infected mice at day 21 pi. This gating strategy discriminates CD4^+^ CD44^hi^ FOXP3^+^ PD-1^+^ CXCR5^+^ T follicular regulatory (Tfr), CD4^+^ CD44^hi^ FOXP3^-^ Ly6C^+^ CD162^+^ Th1, and CD4^+^ CD44^hi^ FOXP3^-^ Ly6C^-^ CD162^-^ PD-1^-^ CXCR5^-^ Th2 cells in our genome-wide expression analysis by RNA-sequencing (RNA-seq). Additionally, we included naive CD4^+^ FOXP3^-^ CD44^lo^ CD62L^hi^ T cells in RNA-seq analysis as a control. Multidimensional scaling (MDS) analysis revealed that the overall transcriptomic profile of naive CD4^+^ T cells, LD- and HD-Tfh cells is highly distinct from each other and the two biological replicates from each group are clustered closely, suggesting high similarities between replicates (**Fig 4A**). In comparison to down-regulated genes, the numbers of up-regulated genes such as *Bcl6 and Cd44* (**S3A-S3B Fig)** were higher in both LD- and HD-Tfh cells over naive CD4^+^ T cells (**Fig 4B**), indicating the Tfh cell-associated activated state. Analysis of differentially expressed genes between LD- and HD-Tfh cells revealed 748 genes with higher expression in HD-induced Tfh cells, and 360 genes whose expression was higher in LD-induced Tfh cells (**Fig 4C**). Although there was no observable difference in gene expression of conventional Th-associated master transcriptional regulator and cytokine transcripts such as *Bcl6/Il21, Gata3/Il4* and *Tbx21/Ifng*, HD-induced Tfh cells had increased expression of genes that have been previously shown to be associated with Th2 cells such as *Il4ra* [32] and *Bhlhe41* (encodes for Dec2) [33], as well as Tfh-associated genes such *Pou2af1* (encodes for Bop1) [34] and *Ascl2* [35] (**Fig 4C**). Conversely, the Th1-associated genes such as *Il2ra* [36], *Il12rb2* [37] and *Il18r1* [21] were highly expressed in LD-induced Tfh cells (**Fig 4C**), suggesting the resemblance of Th1-associated transcriptomic profile in LD-Tfh cells.

**Fig 4.**
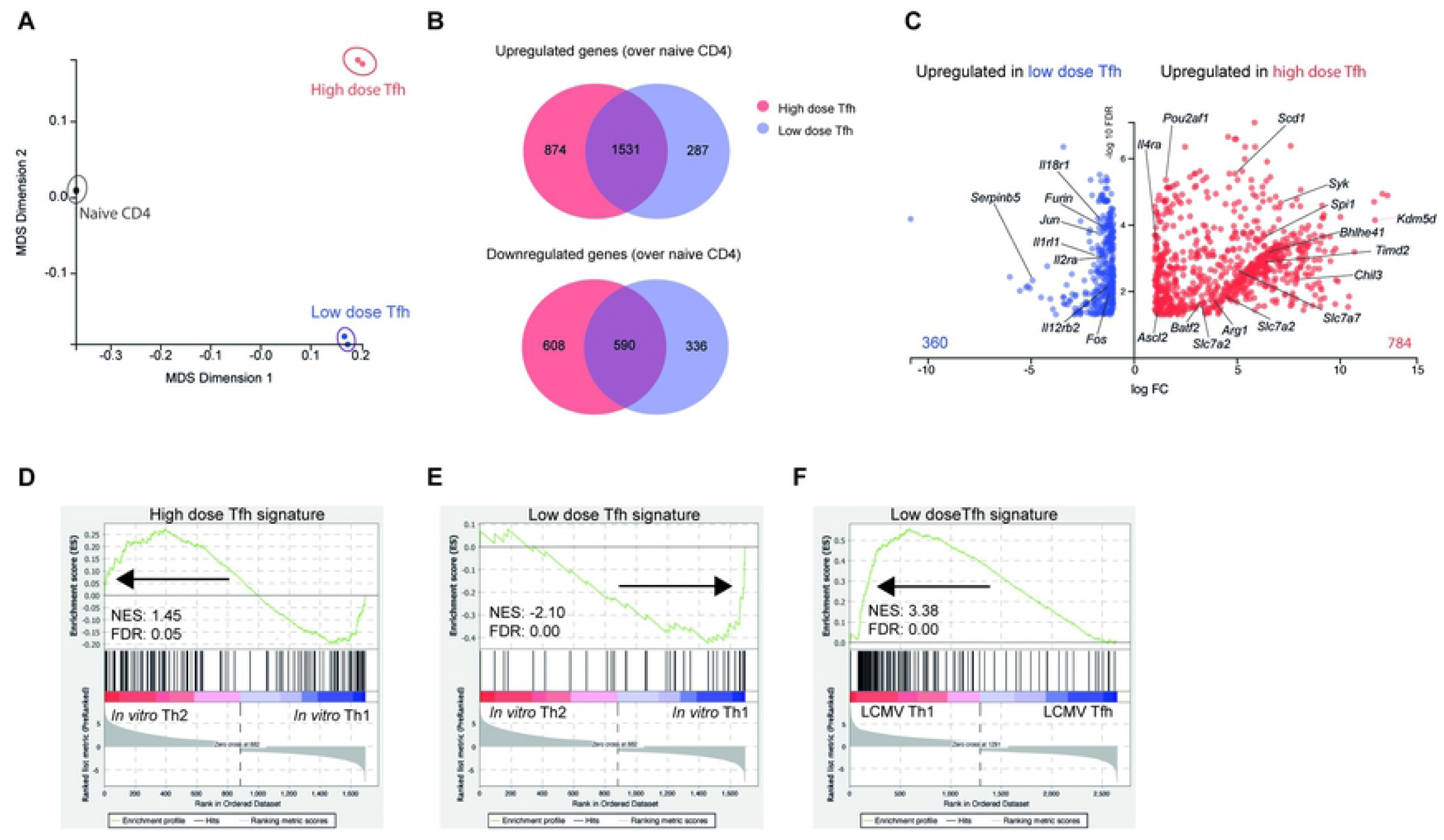
Transcriptional profiles of acute and chronic-induced Tfh cells. FIR mice were infected with acute (high dose) and chronic (low dose) *T. muris*, and mLN-derived Tfh cells (CD4^+^ CD44^hi^ FOXP3^-^ Ly6C^-^ CD162^-^ PD-1^+^ CXCR5^+^) at d21 pi and naive Th cells (CD4^+^ FOXP3^-^ CD44^lo^ CD62L^hi^) from uninfected mice were sorted and purified for RNA-seq analysis. (A) Multidimensional scaling (MDS) analysis of RNA-seq of naive CD4^+^ T cells, low dose and high dose-induced Tfh cells. (B) Number of up- and downregulated genes of Tfh cells/naive CD4^+^ T cells from RNA-seq analysis with > 2-fold (FDR cutoff, 0.05; absolute log-fold > 1). (C) Volcano plot of RNA-seq data showing differentially expressed genes between low- and high-dose-induced Tfh cells with > 2-fold. (D-F) Gene set enrichment analysis (GSEA) of *in vitro* Th1/Th2 cells [39] and LCMV Th1/Tfh cells [7] (F) in high-dose (D) and low-dose (E and F) Tfh gene signatures. Gene expressed >2-fold in high-than low-dose Tfh cells were listed as the high-dose Tfh gene signature and *vice versa* for the low-dose Tfh gene signature. NES, normalised enrichment score; FDR, false discovery rate. Data shown are from one experiment (n=2).

To gain further insights into how similar or different LD/HD-Tfh cells are to bona fide Th1 and Th2 cells, respectively, we first generated LD- and HD-Tfh signature gene sets by listing differentially expressed genes (i.e. ≥ 2-fold) that were highly expressed in LD- or HD-Tfh cells. Using gene set enrichment analysis (GSEA) [38], we examined whether these LD/HD-Tfh gene signatures were enriched in *in vitro* polarised IFN-γ^+^ Th1 and IL-4/13^+^ Th2 cells [39]. As expected, *in vitro* Th2 cells exhibited enriched HD-Tfh but reduced LD-Tfh gene expression signatures (**Fig 4D-4E**), whereas *in vitro* Th1 cells were enriched in LD-Tfh signatures (**Fig 4E**). To better understand whether LD Th1-like Tfh cells are more transcriptionally similar to Th1 or Tfh cells, we also performed GSEA analysis on acute LCMV-induced CXCR5^+^ SLAM^lo^ Tfh and CXCR5^lo^ SLAM^hi^ Th1 cells [7] against our curated LD-Tfh signatures. The analysis further showed a substantial enrichment of the LD-Tfh gene signatures in LCMV-induced Th1 compared to conventional Tfh cells (**Fig 4F**). These observations, taken together, further indicate that LD and HD *T. muris* infection drives Tfh cells that are transcriptionally distinct by sharing Th1 and Th2 cell phenotypes, respectively.

### Th1 versus Th2 cell responses influence chromatin accessibility profile of helminth infection-induced Tfh cells

Given we observed the different transcriptomic profiles of LD- and HD-induced Tfh cells, we next questioned the changes in chromatin accessibility in these cells by performing an assay for transposase-accessible chromatin sequencing (ATAC-seq). In accordance with the transcriptomic data, MDS indicates that the chromatin accessibility of each naive CD4^+^ T cells, LD- and HD-Tfh cells is distinct from each other (**Fig 5A**). We identified over 80,000 accessible genomic regions, where differentially accessible regions in HD-Tfh cells over naive CD4^+^ T cells are higher (i.e., 19,927) when compared to that of LD-Tfh cells (i.e., 16,485) (**Fig 5B**). In contrast, we observed a smaller difference in the numbers of less accessible genomic regions between LD- and HD-Tfh (**Fig 5B**). In addition, we found 7,946 differentially accessible regions between the two groups, in which the majority of these regions (i.e., 5,537) were observed in HD-Tfh cells, while the remaining 2,409 regions were more accessible in LD-Tfh cells (**Fig 5C**). However, these differential accessible regions did not include Tfh-associated locus such as *Bcl6, Il6st* and *Il21r* (**Figs 5C and S4A**). Further, the GSEA analysis showed that LD- and HD-Tfh cell RNA-seq data sets were significantly enriched in ATAC-seq gene signatures, indicating a positive correlation between the transcriptomic profile and epigenetic landscape of Tfh cells (**S4B to S4E Fig**). In support of the enrichment of bona fide *in vitro* Th1/Th2 cell-associated genes in LD/HD-Tfh gene transcriptomic profiles, we found that the RNA-seq data sets of *in vitro* Th1 and Th2 cells were also enriched in LD and HD ATAC-seq gene signatures, respectively (**S4D to S4E Fig**). This enrichment suggests that the chromatin opening across genomic regions in Tfh cells correlates with gene expressions in bona fide Th1 and Th2 cells, depending on the LD versus HD *T. muris* infection, respectively. Gene ontology (GO) analysis using Genomic Regions Enrichment of Annotations Tool (GREAT) [40] revealed that there was a highly significant enrichment of pathways associated with IL-4, IL5 and IL13 secretion and regulation, as well as Th2 cell differentiation in accessible genomic regions of HD-Tfh cells (**S4F Fig**). This prediction further complements transcriptomic profiles that suggest HD infection promotes Th2-like Tfh cell development. In addition, we also observed the GO term negative regulation TLR2 signalling pathway that was enriched in HD-Tfh cells (**S4F Fig**), and given this pathway is critical for Th1 cell differentiation and function [41], this predicts that the absence of TLR2-associated Th1 cell gene expression might in part promote HD-Tfh cell development. In contrast, LD-Tfh-upregulated genes were associated with Th1 cell development and function in response to type 1 pathogens such as viruses, as well as IL-18-induced Th1 cell response that has been previously shown to drive chronic helminth infection [21] (**S4G Fig**).

**Fig 5.**
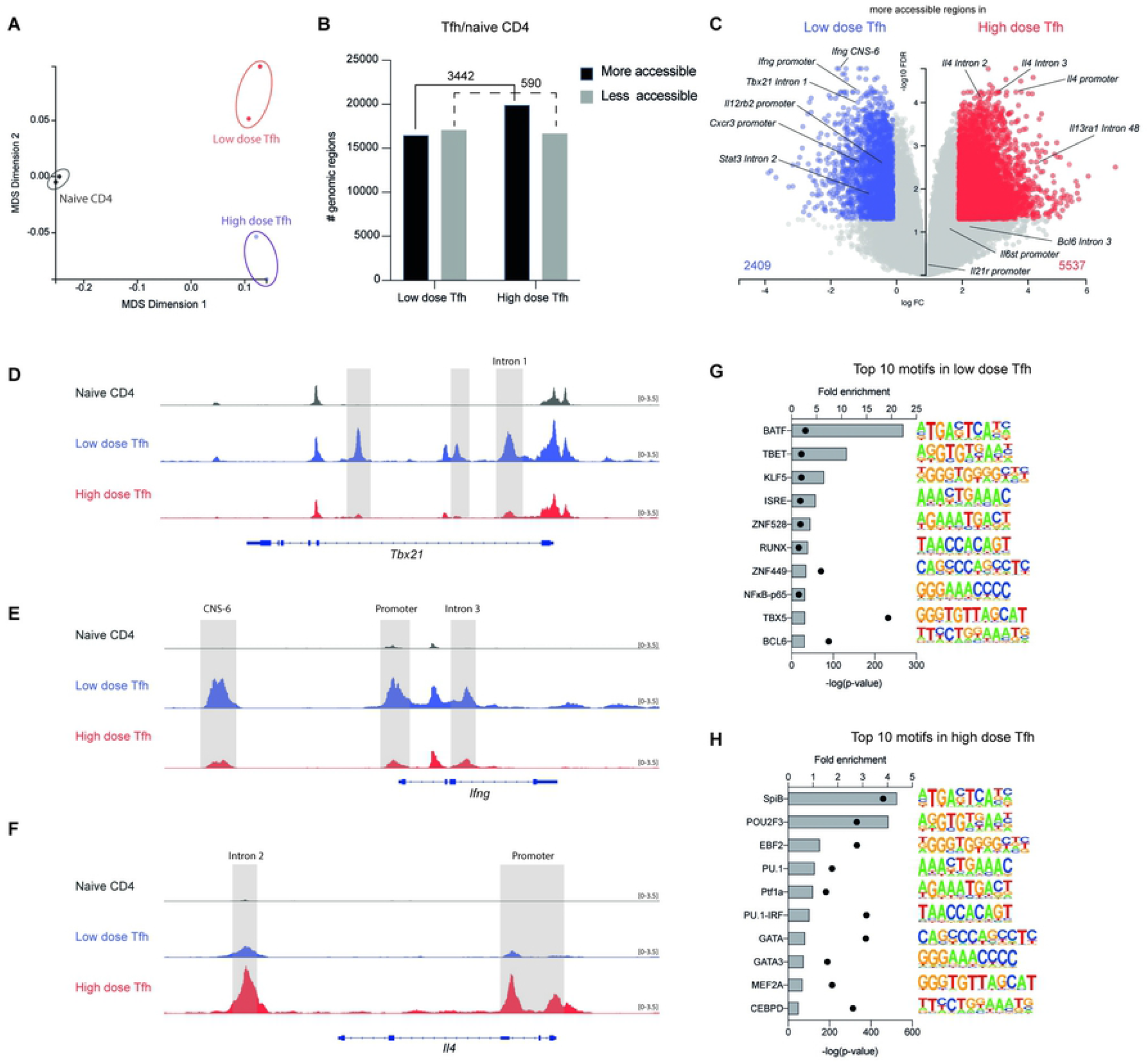
Distinct epigenetic landscape of acute and chronic-induced Tfh cells. Similar gating strategy used for mLN-derived naive Th cells and Tfh cells for ATAC-seq analysis (see Fig 4 legend). (A) Multidimensional scaling (MDS) analysis of ATAC-seq of naive CD4^+^ T cells, low dose and high dose-induced Tfh cells. (B) Number of accessible genomic regions in Tfh cells/naive CD4^+^ T cells from ATAC-seq analysis with > 2-fold (FDR cutoff, 0.05; absolute log-fold > 1). (C) Volcano plot of ATAC-seq data showing differentially expressed accessible regions between low- and high-dose-induced Tfh cells with > 2-fold. (D-F) Integrative Genomics Viewer visualisations of ATAC-seq tracks at the *Tbx21* (D), *Ifng* (E) and *Il4* (F) locus. (G and H) Top 10 transcription factor (TF) binding motif via HOMER de novo analysis in low-dose (G) and high-dose (H) Tfh cells. Graphs show fold enrichment and - log10(p-value) over control ATAC-seq background data. Data shown are from one experiment (n=2).

Given the earlier transcriptomic evidence that suggests Th1- or Th2-like Tfh cells, we next focused on genes related to Th1 and Th2 cells. Despite the higher numbers of accessible regions in HD-Tfh cells (**Fig 5B to 5C**), we found that the chromatin accessibility in Th1-associated genes such as *Tbx21, Ifng and Cxcr3* locus was reduced in HD-Tfh cells compared to Tfh cells from LD-infected mice (**Fig 5C to 5F**). In contrast, the accessibility at *Il4* promoter and enhancer (Intron 2) was increased in HD-Tfh cells (**Fig 5C to 5F**), which is consistent with lower levels of IL-4 expression compared to HD-Tfh cells at d21 pi (**Fig 3B to 3C**). Using TF binding motif HOMER de novo analysis [42], we identified top 10 motifs for LD and HD-Tfh cells that include Th1 and Th2 master TF such as T-bet (**Fig 5G**) and Gata3 (**Fig 5H**), respectively. This TF binding prediction analysis suggests the presence of active T-bet binding activity in LD-Tfh cells that likely binds to accessible *Ifng* locus, leading to IFN-γ expression [43]. In addition, we also found a TF motif for BATF enrichment in LD-Tfh cells (**Fig 5G**). Although BATF is critical for controlling permissive epigenetic modifications within the Th2 cytokine locus and is dispensable for Th1 cell differentiation and function [44], its activity in LD-Tfh cells is likely to be important to promote and maintain Tfh cell development and identity, considering that the absence of BATF leads to the complete loss of Tfh cell generation in response to helminths [44]. Further, the binding motif for Th1-associated TF such as RUNX [45] and p65 [46] (**Fig 5G**) is also enriched in LD-Tfh cells. In contrast, as our analysis predicts Gata3 binding activity in DNA sequences in HD-Tfh cells and given Gata3 directly activates the expression of *Il4* locus [43], this points to the potential regulation of IL-4 expression in HD-Tfh cells by Gata3. Additionally, the analysis also showed the enrichment of TFs such as SpiB, Pou2f3 and PU.1 (**Fig 5H**), with the latter previously implicated in Th2 cell heterogeneity [47] and Tfh cell development [48]. Interestingly, it has been previously shown that Pou2f3 inhibits Gata3 binding activity in Th2 cells [49], and thus pointing to the potential regulation by Pouf3-PU.1 in Tfh cells to regulate Gata3-mediated IL-4 expression. Collectively, these results support the idea that the epigenetic landscape of Th1- and Th2-like Tfh cells are influenced by Th1- and Th2-associated TF binding activity, respectively.

### Th2 cytokines maintain the stability of ex vivo generated Th2-like Tfh cells

As transcriptomic and chromatin landscape analysis suggests that HD-Tfh cells share certain attributes of Th2 cells, this prompted us to assess the stability of HD-induced IL-21^+^ Tfh cells under *in vitro* Th1 and Th2 polarizing conditions. HD-induced IL-21^+^ Tfh cells were sorted at d21 pi, and cultured *ex vivo* for 3 days in either Th1 or Th2 polarizing conditions. We found a higher proportion of cultured cells maintaining the expression of PD-1 and CXCR5 under Th2 than Th1 conditions (**Fig 6A to 6C**). Since intermediate Tfh cells might be a result of partial activation by the polarizing cytokines in the culture system as a transient phase during the course of their differentiation [50], we next assessed IL-4 and IL-21 expression in only non-Tfh and Tfh cells. Most of CXCR5^+^ PD-1^+^ cells expressed higher levels of IL-4 expression under Th2 than Th1 conditions (**Fig 6A, 6B and 6D**), and most of these IL-4^+^ cells co-expressed IL-21 (**Fig 6A, 6B and 6E**). Further, single IL-21^+^ cells were mainly found as CXCR5^+^ PD-1^+^ cells under both Th1 and Th2 conditions, which suggests that IL-21 expression by Tfh cells is not extrinsically influenced by Th1 or Th2 polarising cytokines (**Fig 6A, 6B and 6F**). Unexpectedly, the expanded CXCR5^-^ PD-1^-^ non-Tfh cell population also expressed comparable levels of IL-21-GFP under both Th1 and Th2 conditions (**Fig 6A, 6B and 6F**), suggesting that non-Tfh cells derived from Tfh cells (ex-Tfh) are likely to maintain their ability to produce IL-21 upon activation even in the absence of Tfh polarising conditions. These results indicate that Th2 polarising conditions maintain canonical Tfh cell markers and promote IL-4 expression, highlighting a stability of *ex vivo* generated Th2-like Tfh cells.

**Fig 6.**
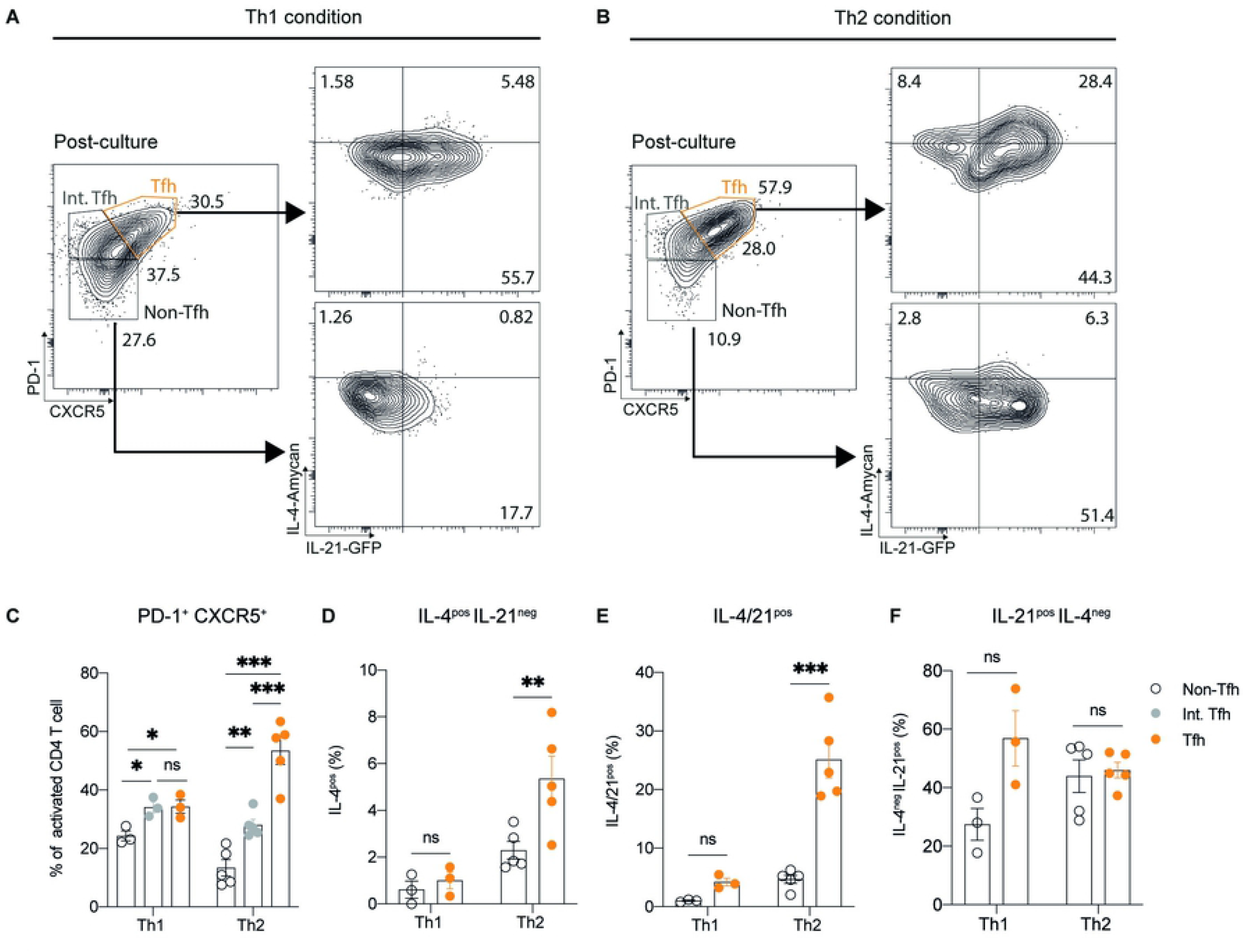
Th2 polarizing cytokines maintain Tfh canonical markers and promote IL-4 expression. (A and B) Representative flow cytometric plots showing IL-4 and IL-21 expression from non-Tfh (CD4^+^ PD-1^-^ CXCR5^-^) and Tfh cells (CD4^+^ PD-1^+^ CXCR5^+^). High-dose-induced d21 pi IL-21-expressing Tfh cells (CD4^+^ PD-1^+^ CXCR5^+^) were cultured under Th1/ Th2 polarizing conditions for 3 days. (C) Frequency of PD-1^+^ CXCR5^+^ expression in cultured non-Tfh, intermediate Tfh (CD4^+^ PD-1^med^ CXCR5^med^) and Tfh cells under Th1/Th2 polarizing conditions. (D-F) Frequency of IL-4^pos^ IL-21^neg^ (D), IL-4/21^pos^ (E) and IL-21^pos^ IL-4^neg^ expression in non-Tfh and Tfh cells under Th1/Th2 polarizing conditions. Data shown are representative of two independent experiments (n = 3–5 mice per group). Statistical significance was determined with a one-way ANOVA. Error bars represent means ± SEM. *p<0.05, **p<0.01, ***p<0.001.

### The loss of T cell-intrinsic T-bet promotes type 2 Tfh-GC responses

Given the importance of T-bet for not only Th1 but also Tfh cell regulation in coordinating the type 1 immune response during viral infection [11–13,15,20], we further characterized whether T-bet plays a role in Tfh cell development in response to helminths. As expected, we observed that HD-Tfh cells expressed higher amounts of Gata3 compared to LD-Tfh cells (**Fig 7A**). As LD *T. muris* infection induces Th1-biased responses, we hypothesised that impaired Th1 cell responses could in turn influence Tfh cell development, and thus the overall Tfh-GC responses. The use of ZsGreen_T-bet [13,51] and IFN-γ_YFP reporter mice revealed that LD-but not HD-Tfh cells expressed T-bet (**Fig 7B**) and higher levels of IFN-γ expression (**Fig 7C**). The expression of IFN-γ in LD-Tfh cells is consistent with a recent finding that uncovers the exclusivity of T-bet expression in Tfh cells in response to the influenza-induced type 1 but not the helminth-induced type 2 immune response [20].

**Fig 7.**
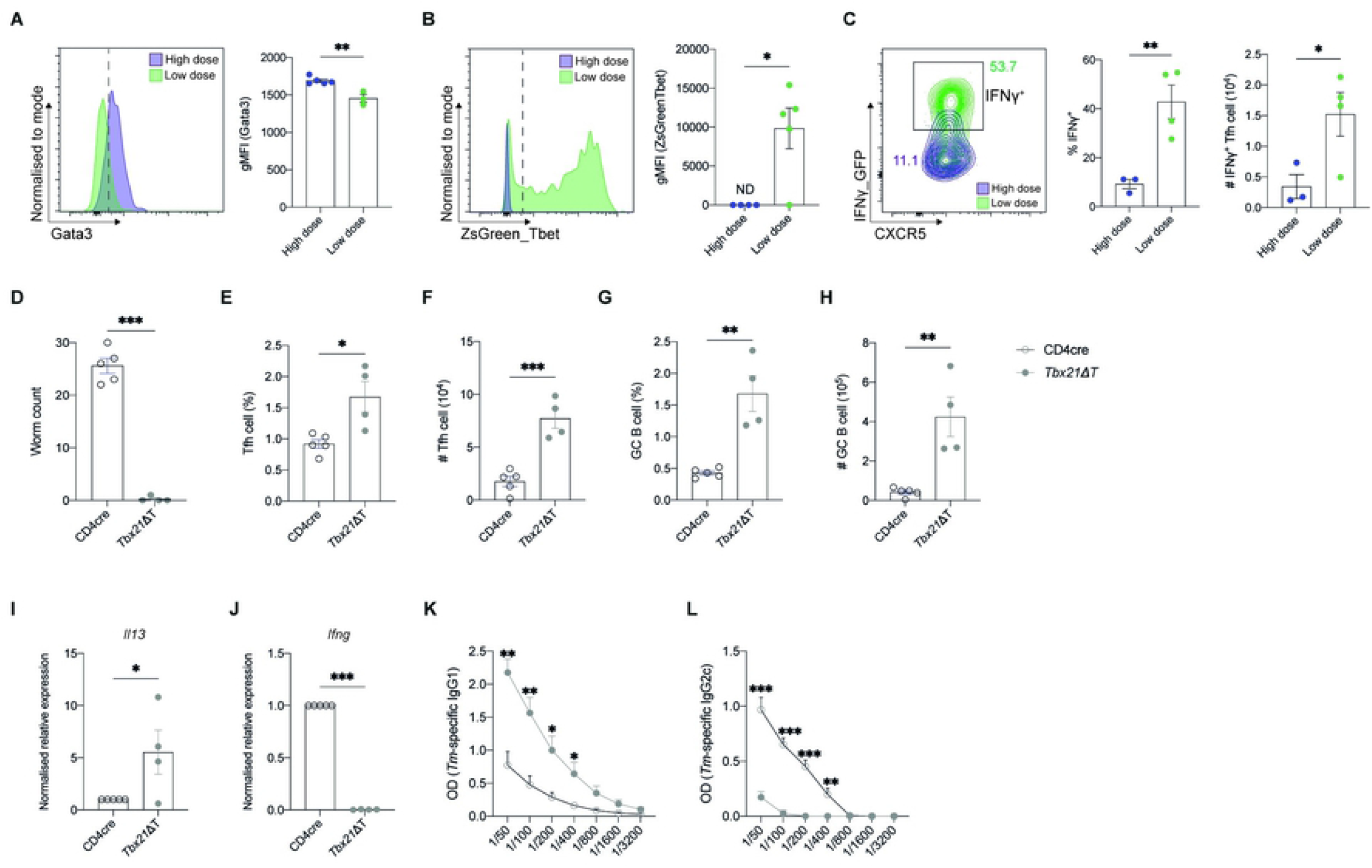
T cell-intrinsic T-bet inhibits Tfh cell development during chronic helminth infection. (A) Gata3 geometric mean fluorescence intensity (gMFI) and frequency in low- and high dose induced-Tfh cells (CD4^+^ CD44^hi^ FOXP3^-^ Ly6C^-^ CD162^-^ PD-1^+^ CXCR5^+^) at d21 pi. (B) ZsGreen_T-bet gMFI and frequency in low- and high dose-induced Tfh cells (CD4^+^ CD44^hi^ FOXP3^-^ Ly6C^-^ CD162^-^ PD-1^+^ CXCR5^+^) at d21 pi. (C) IFNγ_GFP gMFI, frequency and numbers in low- and high-dose induced Tfh cells (CD4^+^ CD44^hi^ FOXP3^-^ Ly6C^-^ CD162^-^ PD-1^+^ CXCR5^+^) at d21 pi. (D) Worm burden analysis in the caecum of low-dose infected Tbx21ΔT and CD4cre control mice at d21 pi. (E and F) Frequency (E) and number (F) of Tfh cells (CD4^+^ CD44^hi^ CXCR5^+^ Bcl6^+^) (F) in mLNs from mice in (D). (G and I) Frequency (G) and number (I) of GC B cells (B220^+^ IgD^-^ CD95^+^ CD38^-^) (F) in mLNs from mice in (D). (J and K) qPCR analysis of *Il13* (J) and *Ifng* (K) of the proximal colon tissue from mice in (D). Numbers shown are normalised expression values to control (CD4cre). (L and M) *T. muris*-specific serum IgG1 (L) and IgG2c (M) from mice in (D). Data shown are representative of two independent experiments (n = 4–5 mice per group). Statistical significance was determined with a two-tailed Student’s t test. Error bars represent means ± SEM. *p<0.05, **p<0.01, ***p<0.001. *Tm*; *Trichuris muris*.

As we had observed T-bet expression in only LD-induced Tfh cells, this prompted us to test an intrinsic requirement of T-bet for Tfh cell expansion following LD *T. muris* infection by using mice specifically lacking T-bet in their CD4^+^ T cells (*Tbx21*ΔT). Additionally, this genetic lesion approach allowed us to examine the effect of a shifted Th1-towards Th2-biased response [52] upon Tfh cell development in response to Th1-biased LD *T. muris* infection. Indeed, *Tbx21*ΔT mice were resistant to LD infection (**Fig 7D**). Importantly, we observed that the loss of T-bet rescued the lack of Tfh cells (**Fig 7E and 7F**) and GC B cell expansion (**Fig 7G and 7H**), which is associated with increased *Il13* (**Fig 7I**) but a concomitant decrease in a non-protective Th1 cytokine *Ifng* (**Fig 7J**) in the proximal colon tissue. As expected, LD-infected *Tbx21*ΔT mice had increased parasite-specific IgG1 but reduced levels IgG2c (**Fig 7K and 7L**), which was also consistent with the prior observation in intact HD-infected C57BL/6J mice (**Fig 2E and 2F**). These results demonstrate that T-bet not only inhibits Th2 cell development [52,53], but also the Tfh cell lineage pathway during type 1-induced chronic helminth infection.

### Diminished GC-Tfh responses promotes susceptibility to acute helminth infection

Although antibodies are not an absolute requirement for protection as their roles are highly context-dependent [2], C57BL/6J mice susceptible to *T. muris* infection in the absence of B cells harbour low levels of Th2 cytokines and concomitantly produce increased levels of IFN-γ [54]. These findings suggest B cells mediate the balance of non-protective type 1 and protective type 2 responses against helminths. Given robust Tfh-GC responses during HD but not LD infection and that HD-Tfh cells share key attributes of Th2 cells, we next questioned whether Th2-like Tfh-GC B cell interaction is required for type 2 protective response. We used anti-CD40L (aCD40L) Ab to disrupt the interaction between Tfh and GC B cells [55] (**Fig 8A**). Worm burden analysis revealed that aCD40L Ab-treated mice cleared most of the parasites; however, there remained low levels of worm burden that had not been fully resolved by d21 pi (**Fig 8B**). We further checked worm burden at d35 pi, a time point by which parasites developmentally moult into an adult stage, and thus is typically used to evaluate complete worm clearance. Similar to LD-infected mice, the parasites remained present in the caecum of aCD40L Ab-treated but not control mice (**Fig 8C**), suggesting that aCD40L Ab treatment compromises protection to the parasite. Immune response analysis in the mLNs showed that the disruption of Tfh-GC responses via aCD40L Ab was successful, as evidenced by impaired Tfh cell development (**Fig 8D-8E**). In addition, we showed that blockade of Tfh-GC interactions resulted in fewer Gata3 expressing Tfh cells (**Fig 8F-8G**), suggesting that cognate B cell-mediated CD40-CD40L signals are indispensable for Gata3 expression in Th2-like Tfh cells. Consistent with reduced Tfh cell numbers, activated B cell (**Fig 8H to 8I**), and GC B cell development (**Fig 8J-8K**) were also impaired upon aCD40L Ab treatment. Given dendritic cells activate naive CD4^+^ T cells through CD40-CD40L interactions to promote Th1 cell differentiation via IL-12 production [56], we wondered if the susceptibility of aCD40L Ab-treated mice was rather due to impaired CD4^+^ T cell activation and Th1 cell response. However, we observed no difference in the levels of CD44 expression in CD4^+^ T cells (**S5A-S5B Fig**), as well as Th1 cell generation (**S5C and S5D Fig**) following aCD40L Ab treatment. Unlike HD-infected mice that generated a strong Th2 cell response, aCD40L Ab treatment during HD infection did not impair Th2 cell development (**S5E and S5F Fig**). Even IgG1 expression amongst GC B cells was not impacted by aCD40L-induced disrupted Tfh-GC responses (**Fig 8L**), the numbers of IgG1-switched GC B cells were reduced (**Fig 8M**) due to the decreased numbers of GC B cells (**Fig 8K**). Further, we showed that levels of parasite-specific IgG1 (**Fig 8N**), and surprisingly, IgG2c were reduced upon aCD40L treatment (**Fig 8O**), despite no difference in plasma cell numbers (**S5G and S5H Fig**). In support of the idea that Tfh-GC responses regulate the type 2 protective response, we found that defective GCs also led to decreased *Il13* (**Fig 8P**) but increased *Ifng* expression **(Fig 8Q**) in the proximal colon tissue, implying that Tfh-GC responses in mLNs indirectly regulate the quality of protective responses at the effector site. Thus, these results demonstrate that Tfh-GC responses are a necessary component of the type 2 protective-dependent immunity to helminth infection.

**Fig 8.**
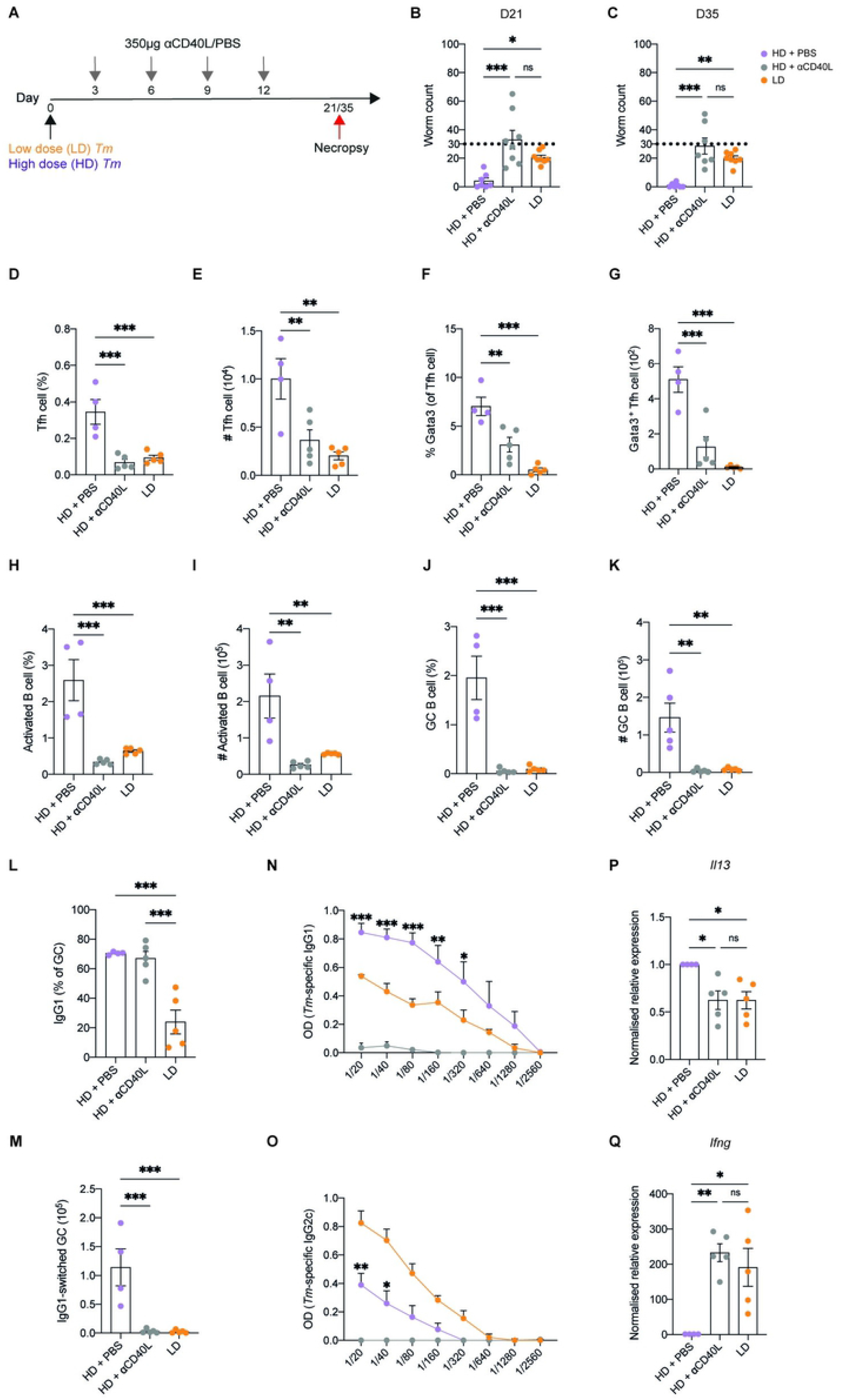
aCD40L Ab treatments impairs Th2-like Tfh and GC responses during acute helminth infection. (A) C57BL/6J mice were infected with acute (high dose) *T. muris* and were treated with 350μg aCD40L Ab (or PBS as a negative control) at d3, d6, d9, and d12, and were analysed at d21 and d35 pi. Low-dose infected mice were included as an additional control. (B and C) Worm burden analysis in the caecum of mice in (A) at d21 (B) and d35 (C) pi. (D and E) Frequency (D) and number (E) of Tfh cells (CD4^+^ CD44^hi^ FOXP3^-^ Ly6C^-^ CD162^-^ PD-1^+^ CXCR5^+^) from mice in (B). (F and G) Frequency (F) of Gata3 expression in Tfh cells (CD4^+^ CD44^hi^ FOXP3^-^ Ly6C^-^ CD162^-^ PD-1^+^ CXCR5^+^) and number (B) of Gata3^+^ Tfh cells from mice in (B). (H and I) Frequency (H) and number (I) of activated B cells (B220^+^ IgD^-^) from mice in (B). (J and K) Frequency (J) and number (K) of GC B cells (B220^+^ IgD^-^ CD95^+^ CD38^-^) from mice in (B). (L and M) Frequency (L) and number (M) of IgG1-switched GC B cells from mice in (B). (M and O) *T. muris*-specific serum IgG1 (N) and IgG2c (O) from mice in (B). (P and Q) qPCR analysis of *Il13* (P) and *Ifng* (Q) of the proximal colon tissue from mice in (B). Numbers shown are normalised expression values to control (high dose + PBS). Data shown are representative of three independent experiments (n = 4–8 mice per group). Statistical significance was determined with a one-way ANOVA. Error bars represent means ± SEM. *p<0.05, **p<0.01, ***p<0.001. *Tm*; *Trichuris muris*.

## Discussion

In this study, we determined a previously unappreciated role of heterogeneous Tfh cell populations that can be used to predict the quality of type 2 protective response to helminth infection. Although Tfh cells are rare during chronic (LD) helminth infection, they have a distinct functional capacity by sharing key attributes of Th1 cells through the expression of T-bet and IFN-γ. In contrast, Tfh cells that develop during a protective Th2-biased response mimic their Th2 cell counterparts by expressing Gata3 and IL-4. Unlike Th1-like Tfh cells that develop during the non-protective Th1 response, the development of Th2-like Tfh cells during acute (HD) helminth infection is consistent with robust GC reactions, extending and confirming previously published studies [16–18,31]. Consistent with their distinct functional capacity, our findings also provide comprehensive transcriptomic and epigenetic profiles of both Tfh cell populations that suggest they are indeed transcriptionally and epigenetically different in Th1 and Th2 phenotypes.

Despite HD-Tfh cells expressing higher amounts of IL-4 than LD-Tfh cells at the peak of Tfh-GC responses (d21 pi), we unexpectedly observed that both Tfh cell populations had a similar capacity to express IL-21 throughout the course of infection. This is consistent with RNA-seq analysis that shows no observable difference between both Tfh cells in the expression of *Icos* and *Klf2*, both of which are critical for regulating IL-21 expression in Tfh cells [7,57]. These findings suggest that while both Tfh cells differ in terms of Th1/Th2 cell attributes, their genetic network that regulates IL-21 expression remains unchanged. In response to viral infection, T cell-intrinsic T-bet is dispensable for IL-21 expression in Tfh cells [12,13], although some studies suggest the opposite [11,25]. Although we did not assess the role of T cell-intrinsic T-bet on IL-21 expression in Tfh cells, we showed that LD-but not HD-Tfh cells express T-bet. However, whether T-bet regulates IL-21 expression in Th1-like Tfh cells in response to chronic helminth infection remains unknown.

Gata3 is critical for IL-4 expression [58], and this is consistent with our data that shows HD-Tfh cells express higher amounts of Gata3 and IL-4 than that of LD-Tfh cells. Further, our work demonstrates that Tfh cells are indeed the major sources of IL-4 in draining mLNs. This finding supports the current dogma that IL-4 secreted by Tfh cells is not only critical for GC formation and IgG1 class switching, but also provides IL-4Rα-mediated signals that regulate the reorganization of stromal cells in LNs during the type 2 immune response [59]. However, given Bcl6 antagonises Gata3 [5,7], the regulatory mechanism that regulates the co-expression of these two TFs, which ultimately allows the development of Th2-like Tfh cells, remains unresolved. Based on how Bcl6 and T-bet expression in type 1-induced Tfh cells are regulated [11], we speculate that Gata3 expression in HD-Tfh cells might be expressed only in a transient manner in preparing the *Il4* locus to be in an open conformation for later expression. During the later stages of Tfh cell differentiation, the balance of Gata3 may be outweighed by Bcl6 expression to allow the acquisition of Tfh cell phenotypes. This speculation requires further experiments using fate-mapping tools and CD4^+^ T cell-intrinsic Gata3/Bcl6 double knockout mice.

In addition to IL-12 [60], IL-2 also suppresses Tfh cell differentiation [61]. Therefore, the high levels of *Il2r* and *Il12rb2* expression in LD-Tfh cells might explain the rarity of these cells. IL-12 directly promotes T-bet expression [53,60]. Although LD-Tfh cells are rare, they highly express T-bet that in turns inhibits Tfh cell development. This is supported by our observation that T cell-intrinsic T-bet deletion elevated Tfh and GC responses, which are associated with the type 2 protective response to a Th1-skewed chronic helminth infection. These data are reminiscent of previous results showing that T-bet represses Tfh cell differentiation in response to type 1 immune stimuli such as *Salmonella* [60] and influenza infection, although this is also dependent on the infection-induced host cytokine milieu [13]. Given IL-12 promotes a chronic *T. muris* infection [62], it is tempting to speculate that a strong IL-12-polarised environment promotes Th1 cell differentiation at the expense of LD-Tfh cell development. At the same time, IL-12 might in part promote the expression of T-bet and IFN-γ in LD-Tfh cells, which is consistent with Th1-like Tfh cell polarisation in the presence of IL-12 [63].

The current dogma of distinct Tfh cell regulation depends on the type of immune response that develops following infection. In expanding this dogma, our results showed that a chronic helminth infection promotes Th1-like Tfh cell response. This indicates that the accompanying immune response determines Tfh cell heterogeneity, irrespective of the class of pathogen (virus vs helminth). At least in the model of *T. muris* infection used in the study, the GC-dependent Th2-like Tfh cell response mediates type 2 immunity to the parasite, presumably through IL-4 production. Given LD-Tfh cells express T-bet, whereas HD-Tfh cells express higher levels of Gata3, this suggests that these two TFs may act as a tipping point that controls the balance between Th1 and Th2 phenotypes in Tfh cells. Although we did not directly assess the direct effects of cytokine cues on Tfh cell responses, we speculate that acute helminth infection drives a strong type 2 cytokine milieu that promotes multiple networks of TFs in Tfh cells, including Gata3. In contrast, a chronic helminth infection induces the type 1 cytokine polarised environment that promotes the adoption of Th1 cell attributes in Tfh cells. However, how the expression of Tfh cell-intrinsic Bcl6, T-bet and Gata3 is dynamically altered throughout helminth infection requires further investigation.

In summary, our work has revealed the heterogeneity of Tfh cells driven by Th1/Th2-biased helminth infection, and that a potent type 2 associated Tfh-GC response is associated with protection to the parasite. Using the helminth *T. muris* model, we identified two subpopulations of Tfh cells that are indeed functionally, transcriptionally, and epigenetically distinct in Th1/Th2 cell phenotypes. However, both of these Tfh cell populations are similar in terms of Tfh cell attributes such as IL-21 expression in spite of the Th-biased response. Thus, Tfh cell heterogeneity can be used to predict the quality of type 2 protective response, which can also be further manipulated to explore any potential therapeutic implications in long-term immune interventions for chronic helminth infection, as well as in a broad spectrum of type 2-mediated diseases such as allergy and asthma.

## Material and methods

### Mice

C57BL/6J mice were obtained from the Monash Animal Research Platform (MARP). IL-21-GFP [25], ZsGreen_T-bet [51], μMT and IL-4Rα knockout (IL-4Rα^-/-^) mice were provided by Stephen Nutt, Joanna Groom, David Tarlington and Nicola Harris, respectively. To generate IL-4-AmCyan-IL-13-DsRed-IL-21-GFP reporter mice, we crossed IL-21-GFP with 4C13R mice [64]. To create IL-4-AmCyan-IL-13-DsRed-IFN-γ-YFP reporter mice, we crossed IFN-γ-YFP [26] with 4C13R mice [24]. Tbx21ΔT mice [13] were bred and maintained with *Cd4*-Cre^+^ mice. FOXP3-IRES-mRFP (FIR) mice [65] were crossed with Ly5.1/2 mice to create FirLy5.1/2-FIR mice. All IL-4-AmCyan-IL-13-DsRed-IFN-γ-YFP, IL-4-AmCyan-IL-13-DsRed-IIL-21-GFP reporter, Tbx21ΔT and FirLy5.1/2-FIR mice were bred and housed under specific-pathogen-free conditions. All mice used in the study were maintained on the C57BL/6J background and were 6-12 weeks of age, and both male and female mice were used, with sex- and age-matched. All experiments using the mice were performed at Monash Biomedicine Discovery Institute (BDI), Monash University in accordance with the Monash University Animal Ethics Committee.

### *T. muris* infection

Mice were infected with either ∼30 (low dose/chronic) or ∼200 (high dose/acute) embryonated *T. muris* eggs by oral gavage. The propagation and maintenance of *T. muris* eggs and parasite-specific antigen were carried out as described previously [19]. Briefly, mice were infected at day 0 and were assessed at different timepoints post infection (pi), as indicated. Worm burden was assessed microscopically by directly counting worms found in the ceca of the infected mouse using a dissecting microscope.

### *T. muris*-specific IgG ELISA

Serum samples from *T. muris*-infected mice were assessed by firstly coating 96-well flat-bottom microplate (Nunc) with 50 μl of *T. muris*-specific antigens at 4°C overnight. Plates were then washed with PBS + 0.05% Tween 20., followed by blocking with 10% NCS for 1 h at room temperature (RT). Diluted serum samples were serially added and were incubated for 1 h at RT. Secondary antibodies (anti-IgG1/IgG2c-HRP), diluted 1: 1000 in 3% BSA/washing buffer were added and incubated for 45 min at RT. TMB substrate was added until the reaction was adequately developed, and was stopped by 1 N HCL. The plates were read at 450 nm.

### Expression analysis/qPCR

RNA was extracted and purified from homogenized proximal colon tissue of *T. muris*-infected mice using standard TRIzol protocol. 1 μg of pure RNA was used for cDNA conversion, followed by expression analysis using qPCR on a SYBR green chemistry (Qiagen) using specific primers: *Il5* fwd 5’-GATGAGGCTTCCTGTCCCTACTC-3’; Il5 rev 5’-TCGCCACACTTCTCTTTTTGG-3’, *Il13* fwd 5’-CCTGGCTCTTGCTTGCCTT-3’; *Il13* rev 5’-GGTCTTGTGTGATGTTGCTCA-3’, and *Ifng* fwd 5’-GGATGCATTCATGAGTATTGCC-3’; *Ifng* rev 5’-CCTTTTCCGCTTCCTGAGG-3’. All samples were standardized against *Actb (*fwd 5’-ACTAATGGCAACGAGCGGTTC-3’; rev 5’-GGATGCCACAGGATTCCATACC-3’).

### Flow cytometry

mLN tissues were mechanically disintegrated and filtered through a 70-μm strainer to form single cell suspensions. Cells were stained using described antibodies (Table S1) as previously described [66]. Data was acquired on a BD LSR Fortessa X20 or LSRIIa., and was analysed using Flowjo (Treestar). Where indicated, cells were sorted and purified by BD FACS Influx.

### Cell culture

CD4^+^ T cells were isolated from the mesenteric lymph nodes (mLNs) of *T. muris*-high dose-infected FIR mice (at day 21 pi) using the StemCell Technologies Inc. EasySep Mouse CD4^+^ T cell Isolation Kit. 1×10^4^ isolated cells were cultured in RPMI media supplemented with 10% heat-inactivated FCS, 100 U/ml penicillin, 100 μg/ml streptomycin, 2mM L-glutamine, 25mM HEPES, and 5×10^−5^ M 2-mercaptoethanol with 1 μg/ml of plate-bound αCD3/28 (clone 145-2c11 and 37.51, respectively). Cells were cultured under Th1- (IL-2, IL-12 [10 ng/ml each] and αIL-4 [10 μg/ml]) and Th2-polarizing conditions (IL-2 [10 ng/ml], IL-4 [40 ng/nl] and αIFN-γ [10 μg/ml]) for 4 days before analysis.

### Generation of mixed bone-marrow chimeric mice

BM cells from μMT and IL-4Rα^-/-^ (or C57BL/6J as a control) mice were mixed in 1:4 ratio to reconstitute lethally irradiated C57BL/6J recipient mice to generate B cell-specific IL-4Rα-deficient BM chimeric mice (B-IL-4Rα^-/-^ mice). 10 weeks after reconstitution, B-IL-4Rα^-/-^ mice were infected with the high dose of *T. muris* eggs and were necropsied at 21 and 35 days later.

### Immunohistochemistry

Tissues from the mLNs were fixed in 10% formalin and paraffin-embedded. 7 μm tissue sections of mLNs were stained with B220 and PNA-biotin antibodies as described previously [67] and Leica ImageScope processing software was used to add 100 μM scale bars.

### Immunofluorescence

mLNs were extracted and were fixed in 4% paraformaldehyde for 6 h and were subsequently immersed in 30% sucrose overnight prior to being embedded in OCT compound. 12-20 μm sections were stained with CD4, GL7 and IgD antibodies as described previously [68]. Images were acquired using a LSM780 confocal microscope (Carl Zeiss MicroImaging) using the acquisition software, Zen Black 2012, as described previously [13]. Images were quantified using image processing software, Image J. 100 μM scale bars were added in all images.

### RNA-seq and bioinformatics

RNA was isolated from FACS-sorted viable CD4^+^ CD44^+^ FOXP3^-^ Ly6C^-^ CD162^-^ CXCR5^-^ PD-1^+^ Tfh cells from the mLNs of high- or low-dose-infected FIR mice after 21 days of infection, using Rneasy Plus Micro Kit (QIAGEN) as per manufacturers’ standard protocol. Viable naive CD4^+^ FOXP3^-^ CD62L^+^ CD44^-^ from the mLNs of uninfected FIR mice were used as a control. RNA was sequenced using Illumina NextSeq 500 (75375, v3) on a MiSeq paired-end run. Raw FASTQ files were analysed with a RNAasik pipeline to produce raw gene count matrix and quality control metrics (Tsyganov et al., 2018). Samples were further analysed with STAR aligner option (Dobin et al., 2013) and reads were quantified with feature counts (Liao et al., 2014) Samples were aligned to the GRCm38 (mm10) transcript. Raw counts were then analysed with a RNA-seq exploration and analysis web-tool, Degust (https://github.com/drpowell/degust), to perform differential expression (DE) analysis and various quality plots (Powell, 2015). DE genes were determined using a FDR cut-off < 0.05 and log2 FC > 1. Data represent the average expression from two biological replicates. Gene enrichment analysis (GSEA) was used to determine the enrichment of published RNA-seq data set (Choi et al., 2020; Scheer et al., 2019) within our curated low- or high-dose Tfh gene signature as previously described (Subramanian et al., 2005).

### ATAC-seq and bioinformatics

DNA was extracted and processed from 5×10^4^ FACS-sorted viable CD4^+^ CD44^+^ FOXP3^-^ Ly6C^-^ CD162^-^ CXCR5^-^ PD-1^+^ Tfh cells from the mLNs of high- or low-dose-infected FIR mice after 21 days of infection, whereas viable naive CD4^+^ FOXP3^-^ CD62L^+^ CD44^-^ from the mLNs of uninfected FIR mice were used as a control, as described by others (Corces et al., 2017; Li et al., 2021). Briefly, cell pellets were DNAse-digested, followed by cell lysis with a chilled lysis buffer prior to nuclei extraction. Nuclei were extracted by resuspending the samples in 50 μl transposition reaction Tn5 mix (Illumina Nextera DNA Library Kit) (FC121-1030) and were incubated in the thermocycler for 0.5 h at 37°C. Subsequently, transposed DNA was isolated using the MinElute PCR Purification Kit (QIAGEN), followed by x5 PCR cycles using a combination of a PCR primer and an index PCR primer. 5 μl of the partially amplified DNA library was used in qPCR reactions to determine the numbers of PCR cycles require to complete library construction. The amplified DNA was extracted using the MinElute PCR Purification Kit (QIAGEN). The DNA library quality and concentration were assessed by using the Bioanalyzer (Agilent) and the Qubit, respectively. ATAC-seq DNA libraries were run as paired-end on the Illumina NextSeq 2000 at Monash Health Translation Precinct (MHTP) Medical Genomics Facility. Read quality was assessed using FASTQC. Reads were aligned to the GRCm38 (mm10) transcript genome and data analysis was performed using the nfcore/atacseq pipeline (Ewels et al., 2020). Data visualization and exploration were performed on Degust (Powell, 2015) (https://github.com/drpowell/degust). DE genomic regions were determined using the FDR cut-off < 0.05 and log2 FC > 1. BedWig files were visualised using Integrative Genomics Viewer (IGV) (Robinson et al., 2011). TF motif enrichment analysis was carried out using HOMER as previously described (Heinz et al., 2010). Gene ontology (GO) enrichment using a web-tool, GREAT (McLean et al., 2010), was used for the GO term enrichment analysis within our curated low- or high-dose Tfh chromatin signature.

### Quantification and statistical analyses

Statistical significance was determined using unpaired (two-tailed) Student’s test or one-way-ANOVA using GraphPad Prism 9 software (GraphPad Software, La Jolla, CA, USA). All graphs are presented as mean ± standard error mean.

### Data and materials availability

RNA- and ATAC-seq datasets described in this study are available at the National Center for Biotechnology Information. Gene Expression Omnibus (GEO) accession numbers: GSE185992 and GSE186062, respectively. RNA-seq datasets from GEO: GSE123966 and GSE140187 were used for GSEA analysis.

## Acknowledgments

We thank Nicola L. Harris, David Tarlinton and Stephen Nutt for providing IL-4Rα KO, μMT and IL-21-GFP mice, respectively. We also thank the staff at the Monash Animal Research facility for technical assistance with animal care and maintenance, the staff at the Monash Flowcore facility for technical assistance with cell sorting, Deanna Lucas and Kirill Tsyganov from Monash Bioinformatics Platform for technical assistance with RNA-seq and ATAC-seq analysis, and all the members of the Zaph and Good-Jacobson labs for technical and intellectual input.

## Supplementary information captions

**S1. Quantitative analysis of GC numbers & IL-13 flow cytometric analysis**.

(A) Quantitation of GC/ section from histological analysis of mLN tissue from Fig. 2A. (B) Geometric mean fluorescence intensity (gMFI) of IL-13-DsRed expression in non-Tfh and Tfh cells following low- and high-dose *T. muris* infection at indicated days pi. Related to Fig. 3. Statistical significance was determined with a two-tailed Student’s t test. Error bars represent means ± SEM. **p<0.01.

**Fig. S2. B cell-intrinsic IL-4Rα is dispensable for IgG1 synthesis but not immunity to helminths**.

(A) Worm burden analysis in the caecum of high-dose *T. muris*-infected C57BL/6J (WT) and IL-4Rα^-/-^ mice at d21 pi. (B-D) Frequency of Th1 (CD4^+^ CD44^hi^ FOXP3^-^ Ly6C^+^ CD162^+^) (B), Th2 (CD4^+^ CD44^hi^ FOXP3^-^ Ly6C^-^ CD162^-^ PD-1^-^ CXCR5^-^) (C) and Tfh (CD4^+^ CD44^hi^ FOXP3^-^ Ly6C^-^ CD162^-^ PD-1^+^ CXCR5^+^) (D) cells. (E) Frequency of GC B cells (B220^+^ IgD^-^ CD95^+^ CD38^-^) (E) and IgG1 in GC B cells (F) from mice in (A). (F) B cell-intrinsic IL-4Rα knockout chimeric mice (B-IL-4Rα^-/-^) setup. Bone marrow (BM) cells from IL-4Rα^-/-^ and μMT mice were mixed in 1:4 ratio to create B-IL-4Rα^-/-^ mice. 10 weeks after BM cell transfer/irradiation, B-IL-4Rα^-/-^ mice were infected with high-dose *T. muris*, and analyzed at d14, d21, and d35 pi. (H) Geometric mean fluorescence intensity (gMFI) of IL-4Rα expression in B cells (CD19^+^) in B-WT control and B-IL-4Rα^-/-^ mice. (I) Kinetics of worm burden analysis in the caecum of mice from in (G) at indicated days pi. (J-L) Kinetics of Tfh cell (CD4^+^ CD44^hi^ FOXP3^-^ Ly6C^-^ CD162^-^ PD-1^+^ CXCR5^+^) (J), GC B cell (B220^+^ IgD^-^ CD95^+^ CD38^-^) (K) and IgG1 expression (in GC B cell) frequency (L) from mice in (G) at indicated days pi. (M-R) *T. muris*-specific serum IgG1 (M-O) and IgG2c (P-Q) from mice in (B) at indicated days pi. Data shown in are representative of two independent experiments (n = 4–5 mice per group). Statistical significance was determined with a one-way ANOVA. Error bars represent means ± SEM. *p<0.05, **p<0.01, ***p<0.001. *Tm*; *Trichuris muris*.

**Fig. S3. RNA-seq data analysis of Tfh cells/ naive CD4**^**+**^ **T cells**.

(A and B) MA plots for differentially expressed genes between high- (A) and low-dose Tfh cells (B) over naive CD4+ T cell control with > 2-fold (FDR cutoff, 0.05; absolute log-fold > 1). Related to Fig 4.

**Fig. S4. ATAC-seq data analysis of Tfh cells & GSEA/ GO enrichment analysis**.

(A) Integrative Genomics Viewer visualisations of ATAC-seq tracks at Bcl6 locus. (B-E) Gene set enrichment analysis (GSEA) of RNA-seq datasets of low/high dose Tfh cells (see Fig. 4) (B and C) and in vitro Th1/Th2 cells (39) (D and E) in curated ATAC-seq high-dose and low-dose Tfh signatures. Genomic regions accessibility >2-fold in high-than low-dose Tfh cells were listed as the ATAC-seq high-dose Tfh gene signature and vice versa for the ATAC-seq low-dose Tfh gene signature. NES, normalised enrichment score; FDR, false discovery rate. (F and G) Gene ontology (GO) term enrichment via GREAT analysis in ATAC-seq data of high- (F) and low-dose (G) Tfh cells. Data shown are from one experiment (n=2). Related to Fig 5.

**Fig. S5. mLN analysis of activated Th cells, Th1/Th2 cells and plasma cells**.

(A and B) (A and B) Frequency (A) and number (B) of activated Th cells (CD4^+^ CD44^hi^ FOXP3^-^). (C and D) Frequency (C) and number (D) of Th1 cells (CD4^+^ CD44^hi^ FOXP3^-^ Ly6C^+^ CD162^+^). (E and F) Frequency (E) and number (F) of Th2 cells (CD4^+^ CD44^hi^ FOXP3^-^ CXCR5^-^ GATA3^+^). (G and H) Frequency (I) and number (J) of plasma cells (B220^-^ CD138^+^). Data shown are representative of three independent experiments (n = 4–5 mice per group). Statistical significance was determined with a one-way ANOVA. Error bars represent means ± SEM. *p<0.05, **p<0.01, ***p<0.001. mLN, mesenteric lymph node. Related to Fig 7.

**Table S1. Reagents/resources used in the study**

